# Urinary Sodium Wasting and Disrupted Collecting Duct Function in Mice with dRTA-Causing *SLC4A1* Mutations

**DOI:** 10.1101/2024.08.21.608692

**Authors:** Priyanka Mungara, Kristina MacNaughton, AKM Shahid Ullah, Grace Essuman, Forough Chelangarimiyandoab, Rizwan Mumtaz, J. Christopher Hennings, Christian A. Hübner, Dominique Eladari, R. Todd Alexander, Emmanuelle Cordat

## Abstract

Distal renal tubular acidosis (dRTA) results in metabolic acidosis due to impaired urinary acidification and can also result in an unexplained urinary sodium-wasting phenotype. Here, we report the generation and characterization of a novel dRTA mutant mouse line, Ae1 L919X knockin (KI). Homozygous L919X KI mice exhibit typical dRTA features including a reduced ability to acidify urine in response to an acid load. This renal acidification defect was associated with a reduced number of Ae1-positive type A intercalated cells. To assess whether these mice exhibit urinary sodium-wasting as seen in some dRTA patients, homozygous KI L919X and the previously described R607H KI mice were fed a salt-depleted acid diet. In line with human patients, both mouse strains exhibited urinary sodium loss. Additionally, we identified increased expression of tight junction proteins claudin-4 and -10b, suggesting a compensatory paracellular pathway in the loop of Henle. Consistent with data from human patients, L919X KI mice displayed a milder phenotype than R607H KI mice. Our findings reveal that both mouse strains are appropriate models for dRTA with a urinary salt-wasting phenotype and a compensatory up-regulation of the paracellular pathway in the ascending limb of the loop of Henle.

## Introduction

Renal tubular acidosis is a group of disorders in which dysregulation of acid-base homeostasis occurs due to improper bicarbonate reabsorption or non-volatile acid excretion by the nephron ^1–5^. The inherited or acquired distal renal tubular acidosis (dRTA) impairs distal proton secretion and consequently renal acid excretion^2,4,6–9^. In addition to alkaline urine, patients with this disease can present hyperchloremic metabolic acidosis ^2,5–9,11^, hypokalemia^10^, nephrocalcinosis / kidney stones^11^, and renal sodium-wasting^12^ with urinary-concentrating defects^5^. Notably, in some patients with inherited dRTA, correction of the metabolic acidosis does not prevent the renal sodium wasting and impaired renal salt handling, consistent with an ion tubular nephropathy^12^.

dRTA affects the collecting duct (CD), a primary regulator of acid base homeostasis and electrolyte and fluid status, which finetunes the final urine composition^6^. The CD is composed of principal (PCs) and intercalated (ICs) cells. PCs mediate electrogenic salt and water reabsorption via the epithelial sodium channel (ENaC) and aquaporin 2 (AQP2)^13,14^. ICs regulate acid-base homeostasis via type A (A-ICs) and type B (B-IC)^15^. A-ICs secrete protons via an apical vacuolar proton ATPase (V-ATPase) and promote reabsorption of bicarbonate via the basolateral kidney anion exchanger 1 (kAE1). B-ICs facilitate bicarbonate secretion through the apical chloride/bicarbonate exchanger pendrin and reabsorb protons via a basolateral V-ATPase. The B-ICs also contribute to electroneutral salt reabsorption through the coupled activity of pendrin and the sodium-dependent chloride/bicarbonate exchanger (NDCBE)^16–18^. At least another intercalated cell type exists, characterized by apical expression of both pendrin and the V-ATPase, which is called non-A non-B ICs. The role of non-A, non-B cells is unclear.

Inherited dRTA may arise from autosomal dominant or recessive mutations in the *SLC4A1* gene encoding kAE1^19–22^. One such dominant mutation is the AE1 R589H, which is located on the cytosolic side of transmembrane domain 6 in the gate domain of kAE1^23–25^. *In vitro* studies have revealed retention in the endoplasmic reticulum^22,26^ and apical mistargeting^27^ of the mutant protein, and a 20-50 % reduction in its anion exchange activity compared to wild-type (WT) kAE1^22^. However, the mutant protein is able to hetero-oligomerize with WT kAE1^26,28^, thereby resulting in a slightly reduced chloride/bicarbonate exchange at the cell surface. AE1 R901X is another common autosomal dominant mutation, which causes a cytosolic C-terminal tail truncation. In *in vitro* studies, the R901X mutant has normal transport activity, but is retained intracellularly and/or apically mistargeted^26,28–30^, similarly to the R589H mutant. However, i*n vivo* data from transgenic mice^31,32^ and from dRTA patients’ kidney biopsies^33,34^ indicate that dRTA-causing variants of kAE1 decrease the abundance of kAE1-positive ICs, emphasizing the limitations of *in vitro* approaches. The kAe1 R607H knock in (KI) mouse model (orthologous to human dRTA R589H mutation) recapitulates classical metabolic acidosis^31^. In this model, R607H KI/KI mice exhibited proper kAe1 basolateral membrane targeting but decreased staining intensity for both kAe1 and the B1 subunit of V-ATPase, and decreased Ae1 mRNA and protein abundance. In addition, the remaining ICs of these mutant mice were larger in size and accumulated the autophagy marker p62. Together, these results are consistent with a reduced number of kAE1 positive A-ICs in the R607H KI/KI mice^31^.

To date, the R607H KI mouse is the only dominant mouse model available. In the current work, we report the generation and characterization of a second dRTA mutant mouse model, carrying the autosomal dominant L919X KI mutation, orthologous to the dominant R901X dRTA human mutation. This second mouse model exhibits the same inability to acidify urine at baseline as the R607H KI mouse line^31^, but overall has a milder phenotype. Additionally, after salt restriction in acid-loaded mice, we found that both L919X KI/KI and R607H KI/KI mice display a persistent urinary sodium loss as seen in dRTA patients. The objectives of this work were to 1) characterize the kAe1 L919X KI mouse line, 2) determine whether the two mouse models accurately reflect the urinary ion loss seen in dRTA patients, and 3) begin investigating the mechanism causing urinary sodium loss in these kAe1 mutant transgenic mice.

## Concise Methods

### Generation of the L919X KI mice

In Germany, animal experiments were approved by the Thüringer Landesamt für Verbraucherschutz (TLV) under the license 02-016/14. In Canada, animal protocols were approved by the University of Alberta’s Animal Care and Use Committee (AUP #1277) and were in accordance with the National and institutional Animal care guidelines. L919X KI mice were generated by homologous recombination similar to Ae1 R607H KI mice^33^ as described in Supplemental Material.

### Metabolic Cage Experimental Design and Urine and Serum Analysis

R607H or L919X mice and WT littermates were provided a “24 hour Salt Depletion or a “14 day Salt Depletion with Acid Load” and placed in metabolic cages (Tecniplast) as described in Supplemental Material. Body weight, chow and water consumption, urine (under mineral oil), and feces were collected every 24 hours. Urine and serum analysis was completed using i-STAT Chem8+ cartridge chip (Abbott Laboratories) and ion chromatography (Dionex Aquion Ion Chromatography System, Thermo Fisher Scientific Inc.).

### Whole Kidney Tissues Analysis

Immediately after collection, a half kidney was either incubated in RNAlater (ThermoFisher) to use for reverse transcription quantitative PCR or a quarter of kidney was decapsulated and processed for immunoblotting as described in Supplemental Material (**Supplementary Tables 1 & 2**). Alternatively, kidneys retrogradely perfused with PFA (4 %) were flash frozen in liquid nitrogen cooled isopentane, and cryosections processed for Masson-Goldner stain, Von Kossa staining or further processed for immunostaining as detailed in the Supplemental Material.

Detailed methods are listed in the Supplemental Material.

## Results

### Generation of Ae1 L919X Knock in Mice and Confirmation of Decreased numbers of A-ICs

We generated mice carrying the orthologous mouse Ae1 L919X mutation by homologous recombination (**Figure 1 A**). Homozygous mice were viable, developed normally and were fertile. Neither gross kidney malformations nor nephrocalcinosis were observed (**Supplementary Figure 1 A & B**), similar to the Ae1 R607H mutant mice^33^. RT-qPCR analysis on whole kidneys from WT and L919X KI/KI mice at steady state confirmed a significant decrease in Ae1 mRNA abundance in the mutant animals (**Figure 1 B**). Immunostaining for kAe1 in the cortex and medulla of WT, heterozygous, and homozygous L919X KI mice (**Figure 1 C & D**) revealed that similar to WT mice, the kAe1 protein predominantly localized to the basolateral membrane, indicating normal membrane targeting of the mutant protein with no evidence of apical localization (**Figure 1 C**). This finding is also consistent with previous findings for R607H KI mice^31^. Additionally, staining intensity of kAe1 in L919X KI/KI mice was markedly reduced compared to WT and heterozygous L919X KI mice, both in the cortex and the medulla. Conversely, signals for pendrin did not exhibit genotype-dependent differences (**Figure 1 C & D**). Collectively, these results suggest a diminished abundance of kAE1-positive A-ICs in the L919X KI mouse model, consistent with Ae1 R607H KI mice observations.

**Figure 1:**
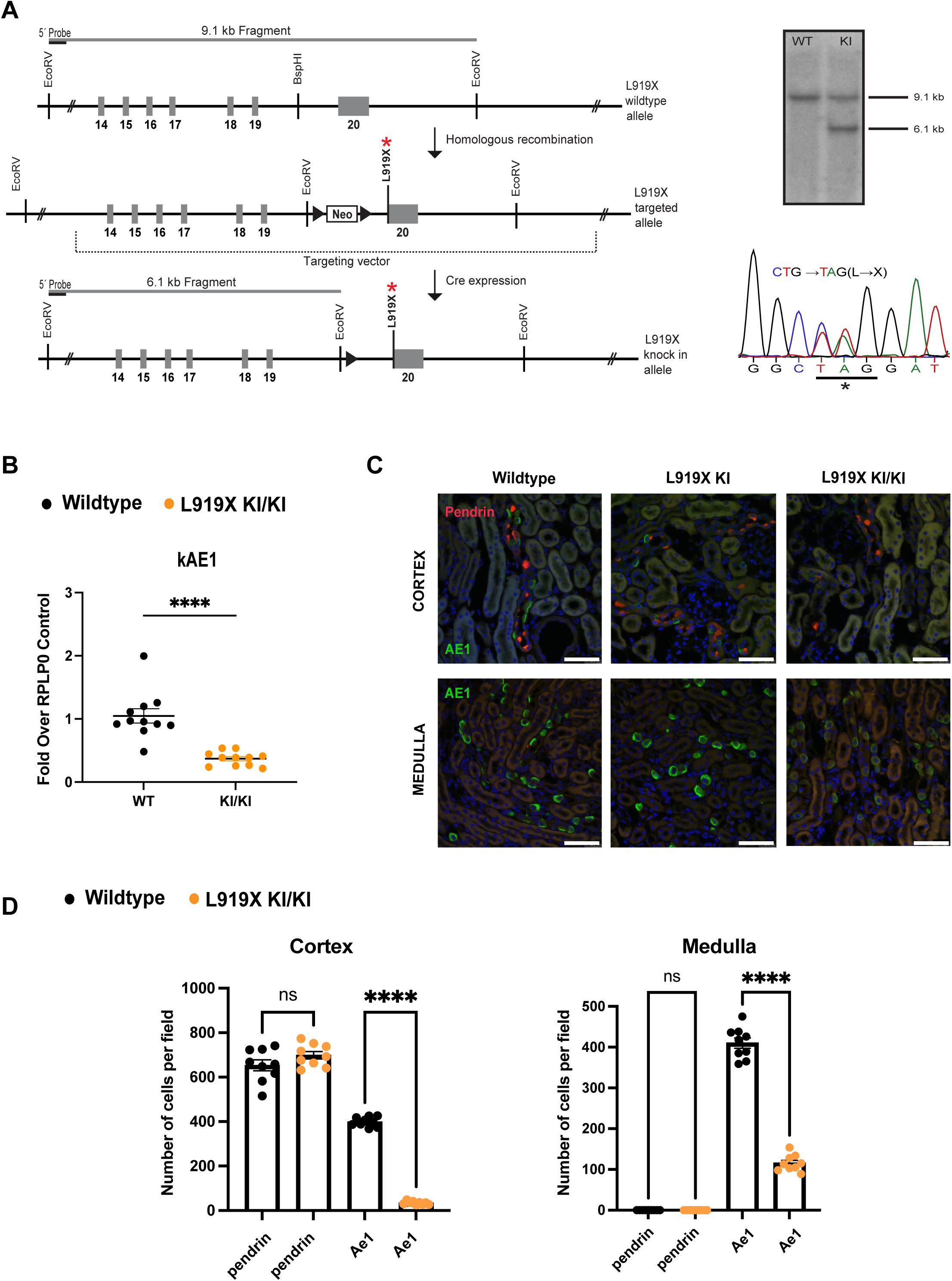
Generation and characterization of the L919X KI mice. (**A**)Targeting and screening strategy with murine Ae1 locus (top) and targeted locus (middle) with locus after removal of the selection cassette by Cre recombination. Right, Southern blot result and sequencing validating the introduction of the mutation to the locus. (**B**) Graph representing the gene expression of Ae1 in perfused whole kidney from WT or L919X KI/KI mice. (C) immunofluorescence images from confocal microscopy of WT, heterozygous or homozygous L919X KI/KI mice medullary or cortical kidney sections stained with anti-pendrin (red) or anti-AE1 (green) antibodies. Scale bar corresponds to 50 µm. (D) Quantification of A-IC (Ae1 positive) and B-IC (pendrin positive) in the cortex and medulla of WT or homozygous Ae1 L919X KI mice. Error bars correspond to means ± SEM, n = 4-6 kidney sections, ***P < 0.001 using Two-way ANOVA.

### L919X KI/KI and R607H KI/KI mice excrete an alkaline urine coupled with less urinary ammonium than WT littermates

To validate the L919X KI/KI mice as a model of human dRTA, we assessed plasma and urine parameters at baseline (**Figure 2, Supplementary Table 3).** L919X KI/KI mice had alkaline urine compared to WT littermates (**Figure 2 A).** In agreement, they also excreted significant less urinary ammonium (**Figure 2 B**). Therefore, the Ae1 L919X KI mice display the characteristic inability to properly acidify urine as seen in dRTA human patients^5^. There was no difference in urine osmolality between genotypes (**Figure 2 C)** and the L919X KI/KI mice excreted sodium, chloride, and potassium comparably to WT mice (**Figure 2 D - F).** There were no differences in plasma ion concentrations or pH in the L919X KI/KI compared to WT mice, although both mouse lines had low plasma bicarbonate and pH, likely due to anesthesia (**Supplementary Table 3).** We completed a baseline analysis of the R607H KI/KI mice as well (**Figure 2 G-L and Supplementary Table 4**). We confirmed that R607H KI/KI mice displayed the same phenotype as L919X KI/KI mice at baseline (**Figure 2 G - L)**. These mutant mice had alkaline urine and decreased ammonium excretion (**Figure 2 G, H),** as previously shown^33^. There was no difference in urine osmolality (**Figure 2 I**) and mutant mice also excreted sodium, chloride, and potassium to a similar extent as WT mice (**Figure 2 J - L)**. Overall, this baseline characterization supports an abnormal urinary acidification in both dRTA mouse lines at baseline, consistent with the hallmark characteristics of human dRTA patients^1^.

**Figure 2:**
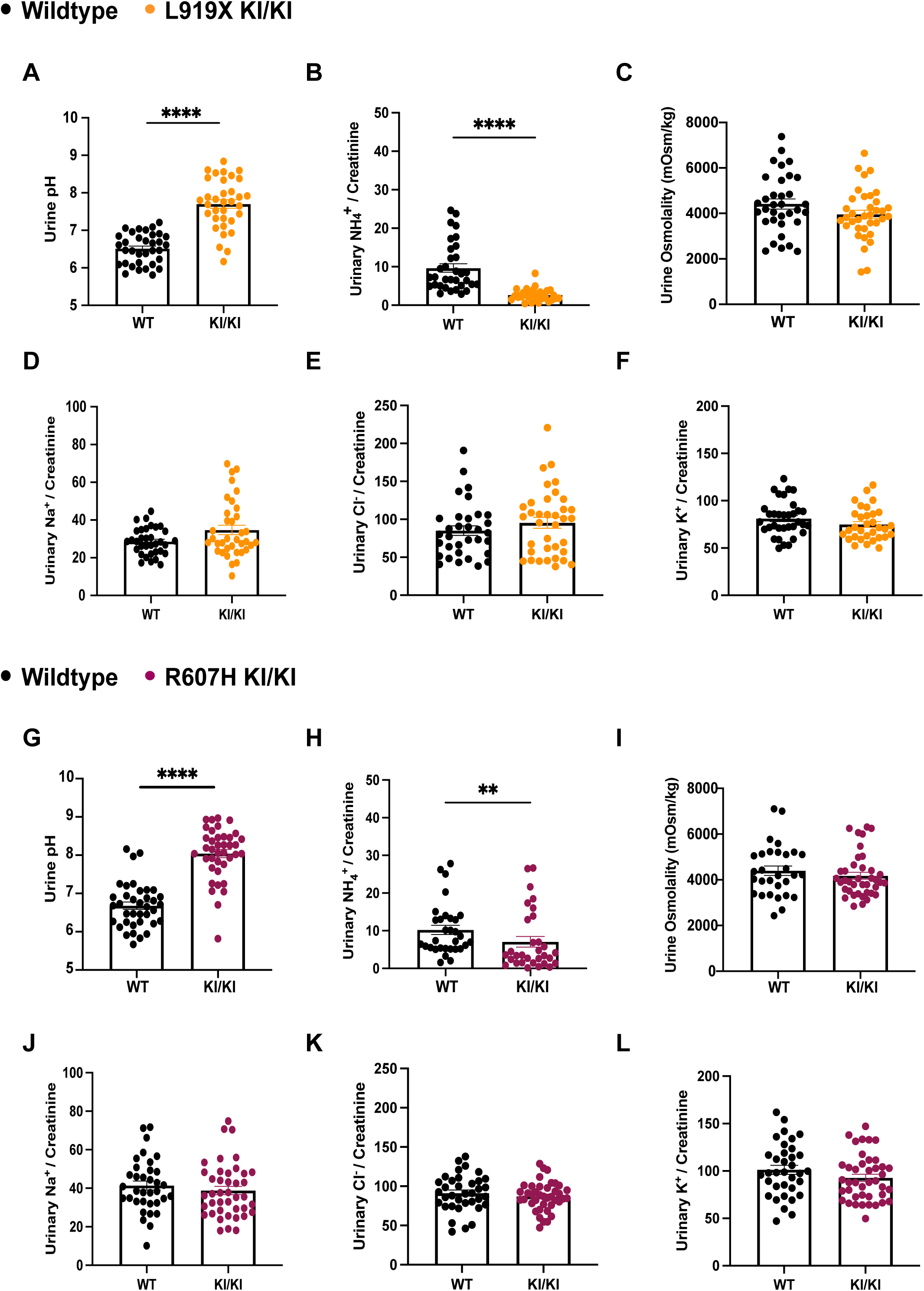
At baseline, both R607H and L919X KI/KI mice exhibit an alkaline urine and decreased urinary ammonium secretion. Urinary pH (**A**), ammonium (**B**), osmolality (**C**), sodium/creatinine ratio (**C**), chloride/creatinine ratio (**D**) and potassium/creatinine ratio (**E**) comparison between WT or L919X KI/KI mice. Urinary pH (**G**), ammonium (**H**), osmolality (**I**), sodium/creatinine ratio (**J)**, chloride/creatinine ratio (**K**) and potassium/creatinine ratio (**L**) comparison between WT or R607H KI/KI mice. Error bars correspond to means ± SEM, **P < 0.01, ****P < 0.0001 using Student’s t-test or Mann-Whitney test.

### Acute sodium depletion confirms impaired urinary acidification and potassium excretion in dRTA mutant mice compared to WT

To assess whether dRTA mutant mice show a salt-losing phenotype similar to dRTA patients, we challenged R607H KI/KI, L919X KI/KI mice or WT littermates with a sodium and chloride-depleted diet for 24 hours (**Protocol 1**) (**Supplementary Figure 2, Supplementary Tables 5 & 6**). Both L919X and R607H KI/KI mice consistently excreted urine with a higher pH compared to WT littermates (**Supplementary Figure 2 A & F**). Both mutant mice strains had no significant differences in urine osmolality between genotypes, although there was trend towards lower osmolality for L919X KI/KI mice (**Supplementary Figure 2 B & G).** Both L919X KI/KI and R607H KI/KI mice reduced urinary sodium and chloride excretion appropriately, to a similar extent as WT littermates (**Supplementary Figure 2 C-D, H-I**). Interestingly, L919X KI/KI mice excreted less urinary potassium while there was only a trend in R607H KI/KI mice (**Supplementary Figure 2 E, J**). No significant plasma difference was found between genotypes for each strain, except that R607H KI/KI mice continued to have increased plasma potassium compared to WT littermates (**Supplementary Tables 5 & 6**). Together these results provide preliminary evidence of renal electrolyte mishandling under acute sodium-depleted conditions for both L919X and R607H KI/KI mice, suggesting a defective coupling of potassium secretion to sodium and chloride reabsorption in the CDs of both strains.

### Salt depleted acid loading results in hypernatremia and hyperchloremia, as well as increased renin levels in both dRTA mutant strains

To further challenge the dRTA mutant mice, we fed WT and mutant mice a sodium-depleted acid diet (Supplementary Material, **Protocol 2**). This diet was designed to recreate a study by Sebastien et al. revealing an urinary sodium wasting phenotype in some dRTA patients on a salt-restricted diet for 8-10 days^12^. Following this challenge, only R607H KI/KI mice displayed metabolic acidosis from the acid load, with decreased plasma bicarbonate and pH (**Supplementary Table 8)**. L919X KI mice only showed a trend for acidosis (p = 0.068) (**Supplementary Table 7)**, suggesting a more severe phenotype in R607H KI/KI than in L919X KI/KI mice. Notably, in both L919X and R607H KI/KI mutant mice, plasma sodium and chloride concentrations were significantly increased compared to WT littermates and compared to steady state (**Figure 3 A & B, G &H, Supplementary Tables 7 & 8**), suggesting a dehydrated state in KI mice. Given the significant hypernatremia and hyperchloremia in both mutant animals, we wondered whether the renin-angiotensin-aldosterone system (RAAS) or dehydration was activated in mutant mice. Indeed, we observed a higher renin mRNA abundance in homozygous R607H KI mice with a similar trend in L919X KI/KI mice (**Figure 3 C, I)**. We next quantified water consumption (**Figure 3 E, K)**, urine osmolality (**Figure 3 D, J**), and plasma hemoglobin and hematocrit levels for both strains (**Supplementary Tables 7 & 8)**. While no difference in urine osmolality was observed compared to WT littermates (**Figure 3 D, J)**, homozygous R607H and L919X mice had reduced water consumption (**Figure 3 E, K)**, despite increased hemoglobin concentration in homozygous R607H mice (**Supplementary Table 8).** These results suggest an abnormal salt handling or impaired osmosensing with activation of the RAAS and/or potential dehydration in these mice.

**Figure 3:**
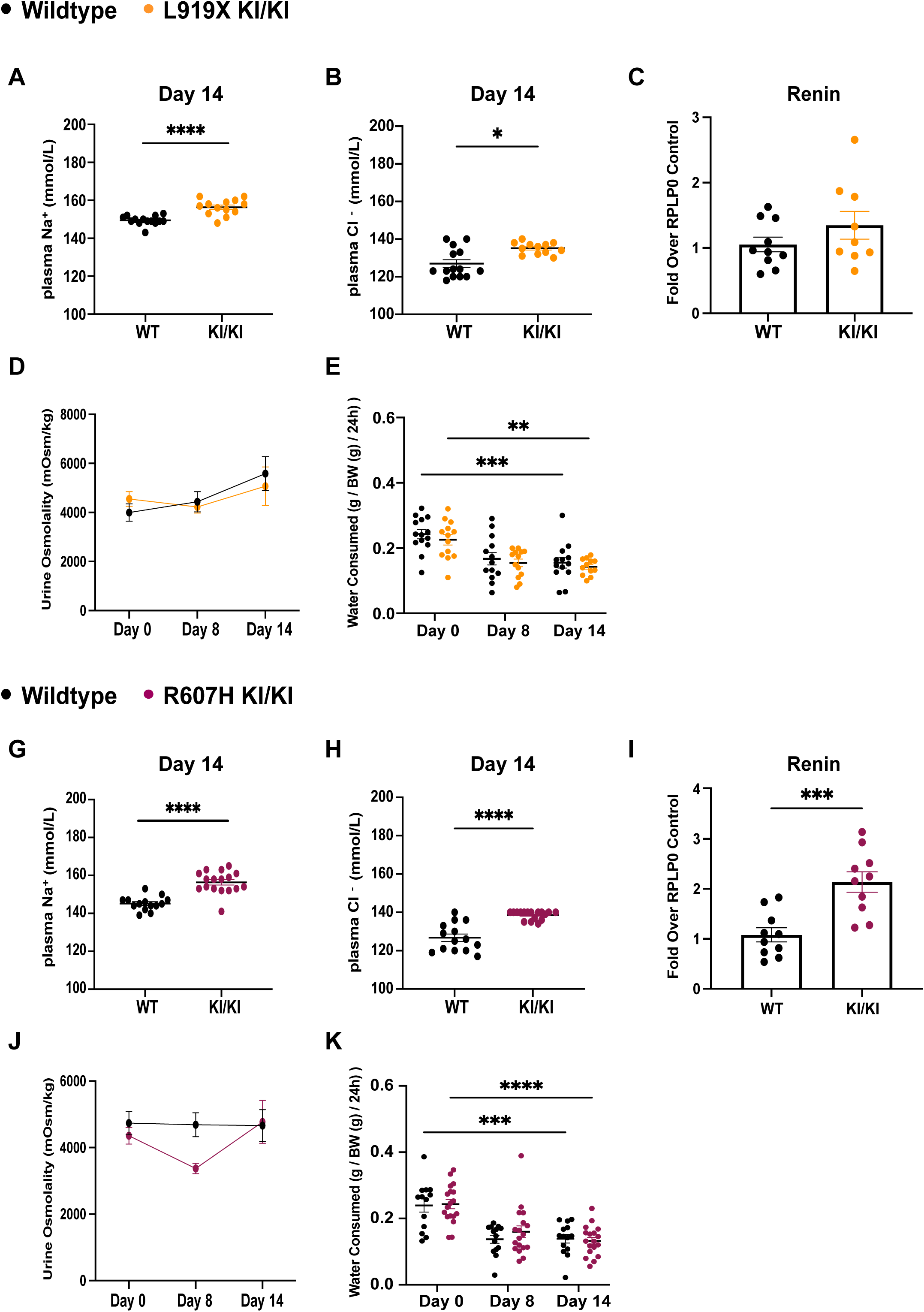
After a salt-depleted acid diet, both R607H and L919X KI/KI mice are hypernatremic and hyperchloremic. Plasma sodium (**A**) and chloride (**B**) concentrations from L919X KI/KI mice or WT littermates. (**C**) Renin gene expression in L919X KI/KI mice or WT littermates. Urine osmolality (**D**) and water consumed (**E)** by L919X KI/KI mice or WT littermates over the 14 day-time course of the diet. Plasma sodium (**G**) and chloride (**H**) concentrations from R607H KI/KI mice or WT littermates. (**I**) Renin gene expression in R607H KI/KI mice or WT littermates. Urine osmolality (**J**) and water consumed (**K)** by R607H KI/KI mice or WT littermates over the 14 day-time course of the diet. Error bars correspond to means ± SEM, *P < 0.05, ***P < 0.001, ****P < 0.0001 using Student’s t-test or two-way ANOVA with Tukey’s multiple comparison test.

### Salt-depleted acid loading reveals a salt wasting nephropathy in homozygous L919X and R607H KI mice

Similarly to the sodium depletion diet alone, both homozygous L919X and R607H KI mice acidified their urine, although not to the same extent as their WT littermates (**Figure 4 A & 5 A).** R607H KI/KI mice excreted significantly less ammonium compared to WT (**Figure 5 B)**, while L919X KI/KI mice (**Figure 4 B)** showed no difference compared to their controls, although both genotypes increased overall ammonium excretion at day 14 compared to day 0 (**Figure 4 B & 5 B).** The salt-depleted acid diet also revealed a pronounced urinary sodium-losing phenotype in homozygous L919X and R607H KI mice (**Figure 4 C&F, Figure 5 C&F**), in agreement with observations from dRTA patients under similar physiological conditions^12^. Distinct phenotypes were observed concerning renal potassium and chloride handling (**Figure 4 & 5 D, E, G, H**). L919X KI/KI mice exhibited urinary chloride loss with no difference in potassium excretion compared to WT mice (**Figure 4 D & E)**. In contrast, R607H KI/KI mice displayed lower urinary potassium and chloride excretion (**Figure 5 D & E)**. These experiments demonstrate that in mice, both the Ae1 L919X and R607H mutation lead to urinary sodium loss and inability to maximally acidify their urine, thus reproducing major features observed in patients suffering with type 1 dRTA.

**Figure 4:**
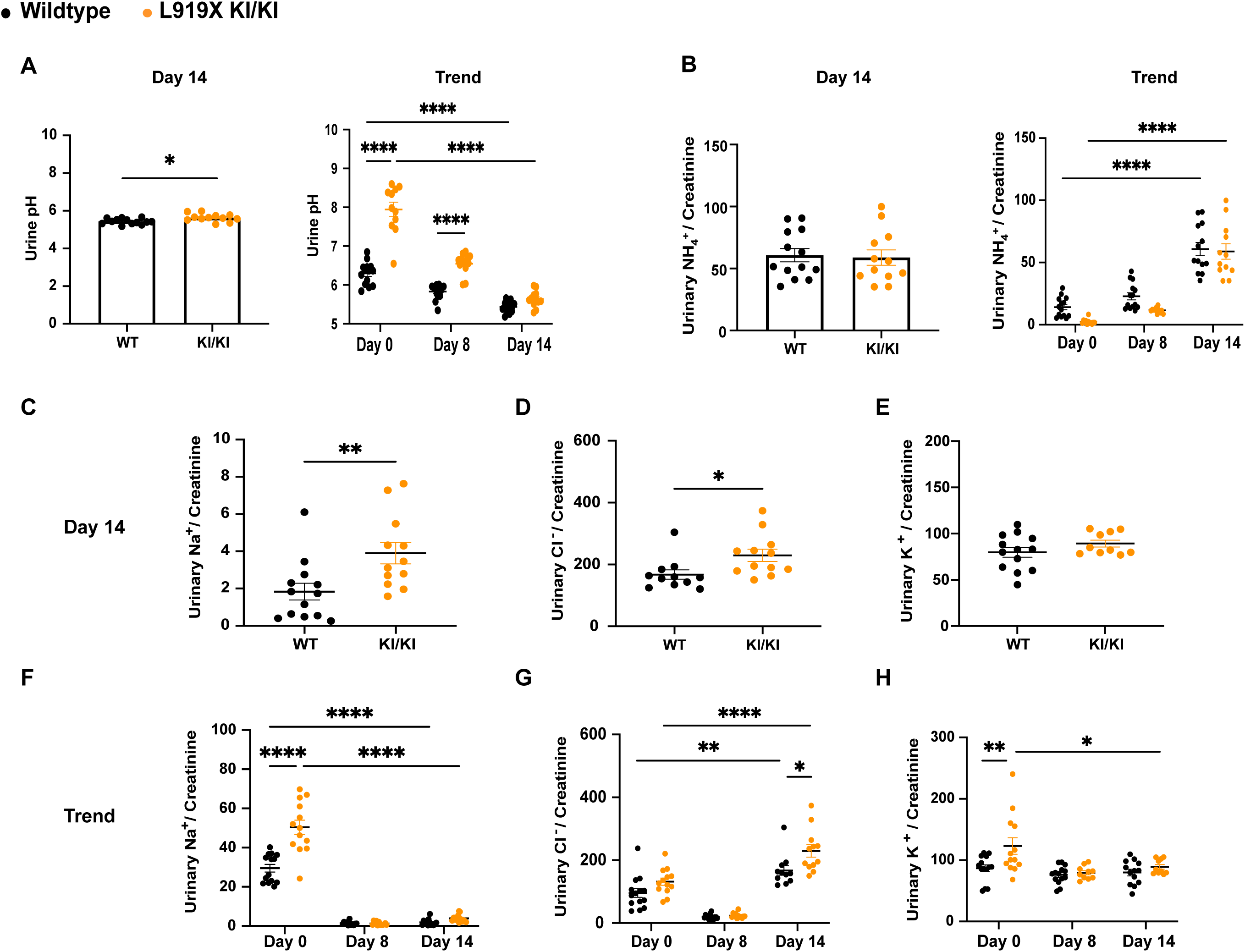
After a salt-depleted acid diet, L919X KI/KI mice produce an alkaline urine and waste urinary sodium and chloride. (**A**) Day 14 (left) and trend (right) over the 14 experimental days urine pH in L919X KI/KI mice or WT littermates. (**B**) Day 14 (left) and trend (right) urinary ammonium/creatinine ratio in L919X KI/KI mice or WT littermates. Day 14 urinary sodium/creatinine ratio (**C**), sodium/chloride ratio (**D**) and potassium/creatinine ratio (**E**). (**F-H**) trend of sodium/creatinine, sodium/chloride and potassium/creatinine ratios over the time course of the experiment in L919X KI/KI mice or WT littermates. Error bars correspond to means ± SEM, *P < 0.05, **P < 0.01, ***P < 0.001, ****P < 0.0001 using Student’s t-test or two-way ANOVA with Tukey’s multiple comparison test. Note that in panel F, no significant difference between WT and KI mice at Day 14 was detected likely due to the two-way ANOVA statistical test comparing multiple conditions and resulting in a high type I error and p value, but a significant difference was found using a Student’s t-test (Figure 3 A).

**Figure 5:**
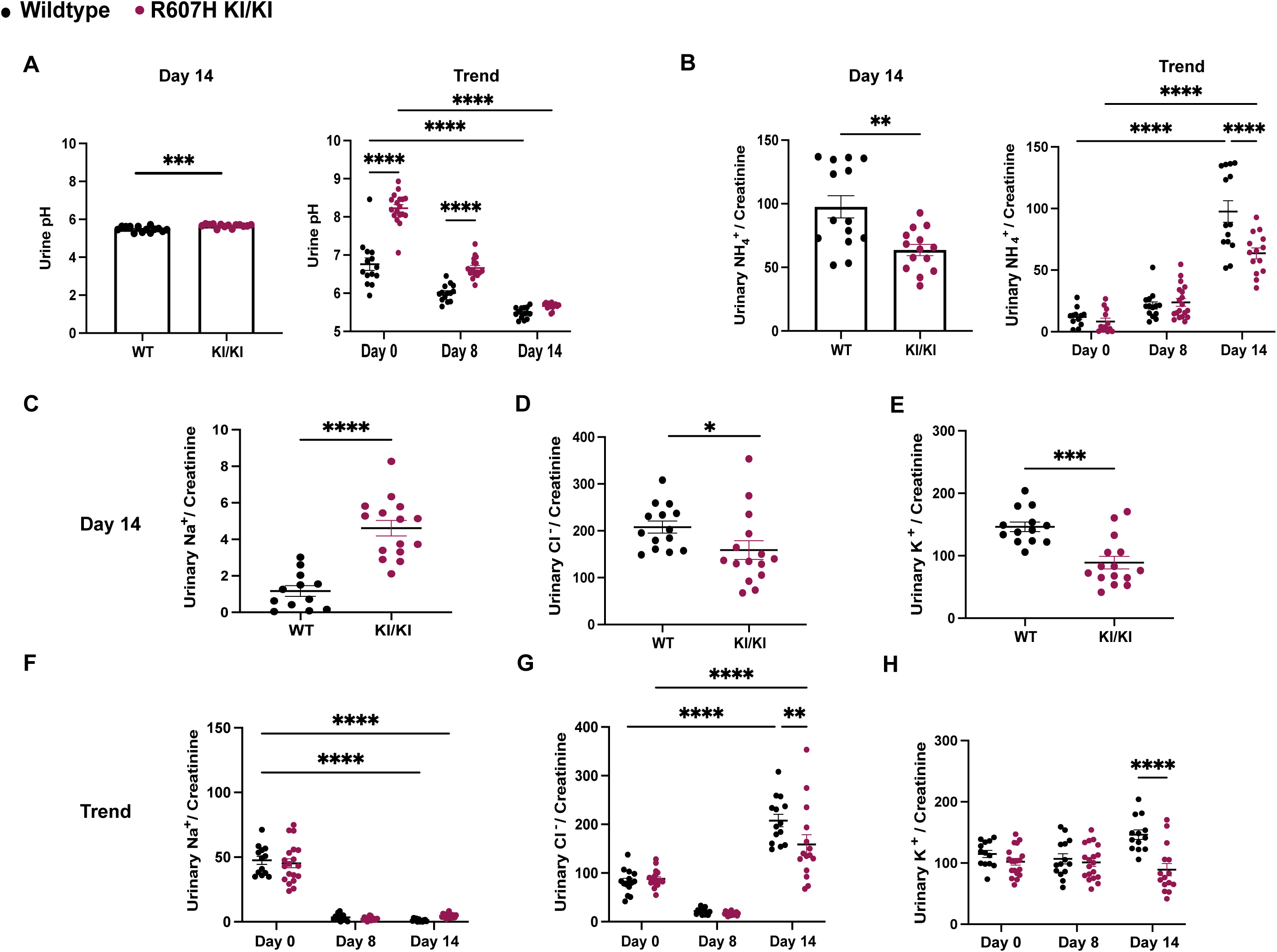
After a salt-depleted acid diet, R607H KI/KI mice produce an alkaline urine and waste urinary sodium and chloride. (**A**) Day 14 (left) and trend (right) over the 14 experimental days urine pH in R607H KI/KI mice or WT littermates. (**B**) Day 14 (left) and trend (right) urinary ammonium/creatinine ratio in R607H KI/KI mice or WT littermates. Day 14 urinary sodium/creatinine ratio (**C**), sodium/chloride ratio (**D**) and potassium/creatinine ratio (**E**). (**F-H**) trend of sodium/creatinine, sodium/chloride and potassium/creatinine ratios over the time course of the experiment in R607H KI/KI mice or WT littermates. Error bars correspond to means ± SEM, *P < 0.05, **P < 0.01, ***P < 0.001, ****P < 0.0001 using Student’s t-test or two-way ANOVA with Tukey’s multiple comparison test. Note that in panel F, no significant difference between WT and KI mice at Day 14 was detected likely due to the two-way ANOVA statistical test comparing multiple conditions and resulting in a high type I error and p value, but a significant difference was found using a Student’s t-test (Figure 3 G).

### Paracellular proteins claudin 4 and claudin 10b are upregulated in dRTA mutant mice

Given the alkaline urine and the urinary sodium loss after salt depleted acid loading, we quantified mRNA and protein levels of specific markers of the CD, the thick ascending limb (TAL) and the distal convoluted tubule (DCT) in KI/KI mice and the respective WT littermates (**Figure 6, Supplementary Figures 3 - 4)**.

**Figure 6:**
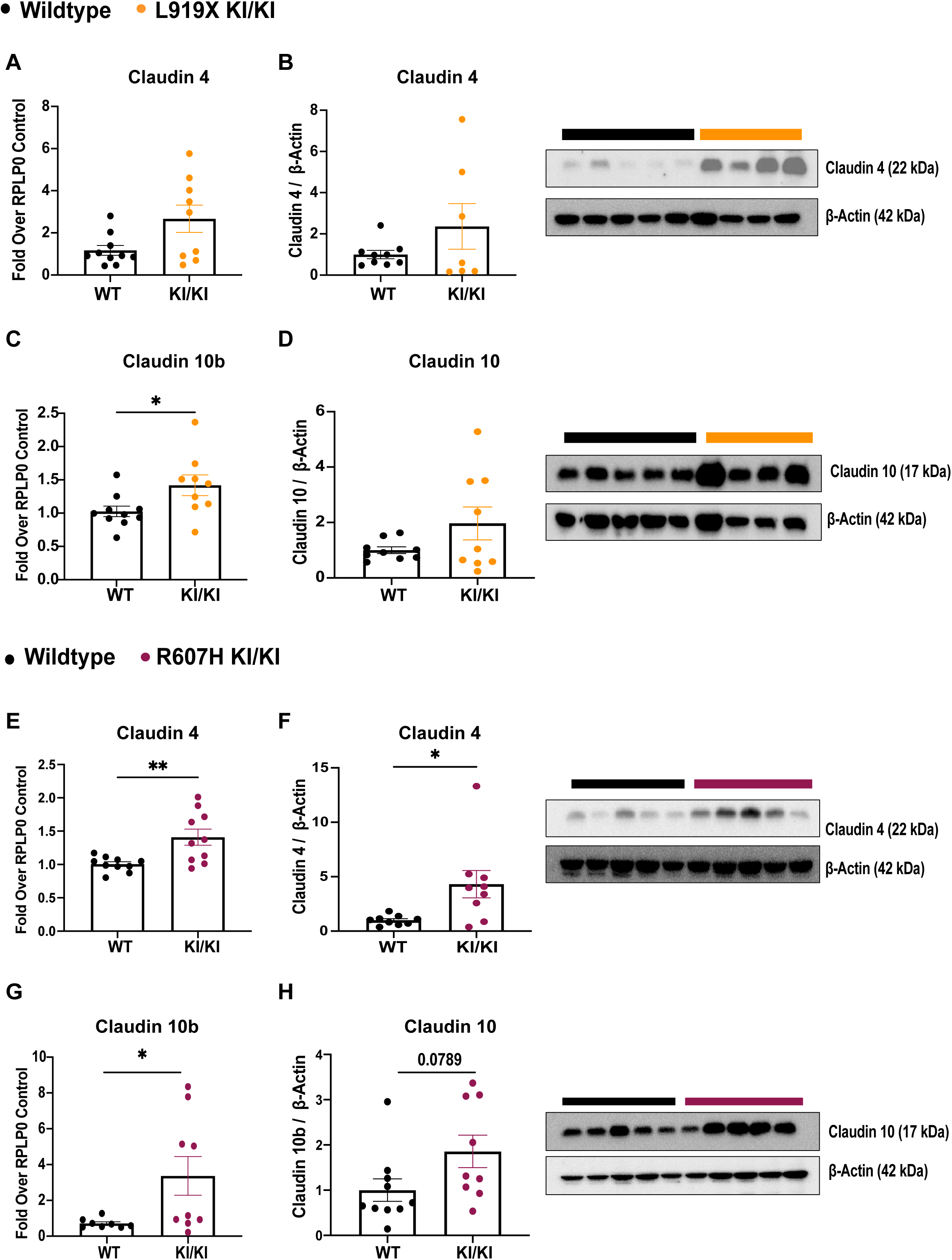
Claudin-4 and claudin-10b are upregulated in both L919X and R607H KI/KI mice after a salt depleted acid diet. Claudin-4 gene expression (**A** & **E**) and protein abundance (**B** & **F**) with a representative immunoblot image (right). Claudin-10b gene expression (**C** & **G**) and claudin-10 protein abundance (**D** & **H**) with a representative immunoblot image (right). Error bars correspond to means ± SEM, *P < 0.05, **P < 0.01, using Student’s t-test or two-way ANOVA with Tukey’s multiple comparison test.

We found increased mRNA and protein levels of the paracellular pores claudin 4 and claudin 10b, in both L919X KI/KI and R607H KI/KI mice (**Figure 6 A-H**). Claudin-4 is a paracellular chloride pore and a sodium pore blocker expressed in the thin ascending limb of Henle’s loop, the distal tubule and CD^35–37^, while claudin 10b is a sodium pore in the TAL^38–40^. These findings suggest a common compensatory mechanism via the paracellular pathway in the loop of Henle of both mutant strains.

The salt-depleted acid challenge also revealed a downregulation of A-IC and B-IC marker gene expression in R607H KI/KI mice, with similar trends in L919X KI/KI mice (**Supplementary** Figure 3 **& 4 A - I**). There was no clear trend for principal cell markers between KI/KI mice and WT littermates. In agreement with decreased numbers of ICs reported for R607H KI/KI mice^31^, kAe1 and B1-ATPase mRNA or protein abundances in whole kidney lysate were significantly reduced compared to WT littermates (**Supplementary Figure 3 & 4 A, C, K**). Messengers for pendrin and AE4 were significantly reduced in R607H KI/KI mice, with a similar trend in L919X KI/KI mice (**Supplementary** Figure 3 **& 4 C, E**). Increased NDCBE mRNA abundance was observed in R607H KI/KI mice with a similar trend in L919X KI/KI mice (**Supplementary Figure 3 & 4 D**).

Given the higher abundance of the TAL-specific claudin-10b in both mutant mouse lines (**Figure 6 C, D, G, H**), we investigated gene expression of various transporters from prior nephron segments (**Supplementary Figure 3 & 4 H – J, M**). The mRNA abundance of the DCT sodium-chloride cotransporter (NCC) was significantly lower in L919X KI/KI mice with a similar trend in R607H KI mice (**Supplementary Figure 3 & 4 J**). We finally assessed the mRNA abundance of the sodium/potassium/chloride cotransporter NKCC2 and sodium/proton exchanger NHE3 in whole kidneys, with the caveat that NHE3 is also abundantly expressed in the proximal tubule. These levels did not differ between L919X KI/KI and WT counterparts, but were significantly decreased in R607H KI/KI mice compared to WT littermates (**Supplementary Figure 3 & 4 H, I**). Overall, these results highlight paracellular adaptations in both KI/KI mice, potentially in an attempt to maintain salt homeostasis.

## Discussion

In this study, we characterized a second dRTA mouse model that carries the murine Ae1 L919X mutation orthologous to human dominant R901X dRTA mutation. We focused on homozygous mice as they display a stronger phenotype than heterozygotes more representative of dRTA patients’ symptoms. In agreement with the previously published R607H KI/KI model, at baseline, the L919X KI/KI mice exhibit an alkaline urine compared to WT littermates, mimicking the main physiological defect observed in dRTA patients. In perfused kidneys from homozygous L919X and R607H KI mice, we observed decreased Ae1 gene expression suggesting a loss of A-ICs. Overall, our results support that both L919X and R607H KI/KI mouse lines reproduce the alkaline urine seen in dRTA and thus are good models to investigate the underlying pathophysiology.

Our results also reveal a more pronounced phenotype in R607H KI/KI compared to L919X KI/KI mice with slight differences in abundance of mRNA of Henle’s loop and CD markers. One study reporting dominant dRTA mutations in the *SLC4A1* gene found that despite phenotypic similarities, the R901X mutation caused a milder phenotype than the R598H. This includes a lower plasma bicarbonate concentration and a more severe hypokalemia in patients with the R589H mutation compared to R901X^21^. Our results confirm this trend with significant differences observed in R607H KI/KI mice versus WT littermates but only similar trends in the L919X KI mice.

We challenged homozygous Ae1 L919X and R607H KI/KI mice and the respective controls with an acute salt-depleted diet or a salt-depleted diet in combination with an acid challenge to assess whether these mice develop a sodium wasting phenotype as shown for some dRTA patients^12^. Both dRTA mutant mice exhibit renal sodium loss compared to WT controls. Both KI/KI mice had elevated plasma sodium and chloride levels with increased abundance of renal renin mRNA suggestive of a volume contracted state despite a similar volume of water consumed and urine osmolality. Importantly, we found an increase in claudin 4 and claudin 10b gene and protein expression in both homozygous L919X and R607H KI mice compared to WT. A previous study feeding a salt restricted diet to a B1-H^+^-ATPase knockout mouse model elucidated one mechanism: loss of *Atp6v1b1* impaired sodium reabsorption via pendrin/NDCBE in B-ICs. These mice also had increased urinary ATP and prostaglandin E2 (PGE2) release (known sodium movement inhibitors), which decreased ENaC expression on PCs further contributing to sodium wasting^41^. Consistent with these findings, our experiments showed decreased pendrin and Ae4 gene expression levels, while NDCBE gene expression was increased in R607H KI/KI mice. We also observed increased γ-ENaC (SCNN1G) mRNA and protein abundance in R607H KI/KI, but no significant difference in renal outer medullary potassium channel ROMK gene expression. These trends were also observed in L919X KI/KI mice, with the additional finding of decreased NCC mRNA abundance.

Both mouse lines displayed a similar alteration in their paracellular pathway in Henle’s loop as an increase in claudin-4^6,45,4^ and claudin-10b^42^ mRNA abundance was observed after acid challenge with salt depletion. Claudin-4 functionally interacts with kAe1, as expressing the exchanger in immortalized inner medullary collecting duct cells increases transepithelial permeability to both sodium and chloride in a mechanism dependent in part on claudin-4 and with-no-lysine kinase 4^42,43^. Although present in tight junction, claudin 10b also colocalizes with the sodium/potassium ATPase and chloride channels in the basolateral infoldings of the TAL^39^. While its role there remains unclear, it may play a crucial role in facilitating high electrolyte flow, and preventing basolateral water influx. Therefore, increased expression of claudin 10b may represent a compensatory mechanism to enhance urine concentration and regulate electrolyte balance.

The upregulation of claudins in Henle’s loop points towards a possible compensatory role of this segment in urinary acidification and sodium reabsorption. The ascending limb of Henle’s loop is a site for urinary acidification via the combined action of apical NHE3 and basolateral NBCn1^44,45^. As shown by our experiments, mutant mice still acidify their urine although they are kAE1-depleted, which points towards a possible contribution of this segment to urinary acidification, in addition to sodium and chloride reabsorption. The function of acid-base transporters is tightly coupled to that of salts in the TAL, through apical NHE3 and the basolateral electroneutral sodium bicarbonate co-transporter, NBCn1^49, 50^. NBCn1 facilitates urine acidification during metabolic acidosis^50^, promoting ammonium reabsorption across the renal tubules, rapidly restoring arterial blood pH and facilitating its secretion by the CD^51^. Previous work reported that R607H KI/KI mice have increased NCBn1 protein abundance at steady state compared to WT mice^33^. The potential upregulation of this protein in the KI/KI mice lines could provide an explanation for their ability to effectively regulate plasma pH and acidify their urine to some extent despite the loss of IC. Additionally, NKCC2 and ROMK in the TAL may cause a net chloride reabsorption creating a lumen-positive transepithelial potential driving paracellular sodium reabsorption through claudin 10b.

After the salt-depleted acid challenge, we detected a significantly decreased mRNA abundance of NHE3 and NKCC2 in homozygous R607H KI mice compared to WT. The relevance of this finding may be limited as both NHE3 and NKCC2 proteins are post-translationally regulated. NKCC2 regulation occurs via membrane trafficking, phosphorylation, and protein-protein interactions^46^. NHE3 is regulated through phosphorylation, and its activity can be inhibited by aldosterone^47,48^, which may be relevant in the context of salt depletion.

There are some limitations to this work. Although the KI/KI mice reproduced some symptoms of dRTA, they are not exhibiting them all. First, these mice knocked in with a dominant dRTA mutation display a milder phenotype than dRTA patients. Indeed homozygous mice exhibited acidemia only after an acid challenge^31^, while heterozygous dRTA patients with dominant mutations have overt acidemia at baseline. We hypothesize that this is due to the difference in diet between rodents and humans. Therefore, we have focused our study on homozygous mice. Second, these mice do not exhibit hypercalciuria, nephrocalcinosis and kidney stones that are commonly observed in dRTA patients. The reason for this discrepancy is unknown. Importantly, mice with a disruption on *Atp6v1b1*, which in humans causes dRTA, also lack the hypercalciuria, nephrocalcinosis and overt acidemia^41^ seen in these patients, pointing to some limitations inherent to these mouse models. However, although a limitation, the lack of these symptoms allowed us to investigate other dRTA symptoms without the caveats of kidney stones and nephrocalcinosis.

Overall, our results show that both L919X and R607H and KI/KI mice recapitulate some dRTA symptoms, including the urinary sodium loss seen in patients. Our results reveal physiological mechanisms for this complex disease and point to a potential compensatory acidification and salt reabsorption in earlier nephron segments.

## Acknowledgments

This study was funded by the Canadian Institutes of Health Research (PJT#168871) and the Kidney Foundation of Canada (2020KHRG-666615) to E.C. C.A.H. was supported by a grant of the DFG (HU 800/7-2). D.E. is supported by a grant from the French National Research Agency (ANR-14-CE12-0013-01), and by the National Center for Precision Diabetic Medicine – PreciDIAB. (ANR-18-IBHU-0001). P.M. received a Canada Graduate Scholarship-Master’s graduate studentship from the Canadian Institutes of Health Research, a University of Alberta Graduate Recruitment Scholarship and a Walter H. Johns Graduate Fellowship from the University of Alberta. A.K.M.S.U. received an NSERC CREATE graduate studentship, G.E. received a Graduate Student Engagement Scholarship, a Faculty of Medicine and Dentistry Delnor Scholarship and a Faculty of Medicine and Dentistry 75^th^ Anniversary award. F. C. was awarded an University of Alberta Graduate Recruitment Scholarship.

---

**Supplementary Table 1:**
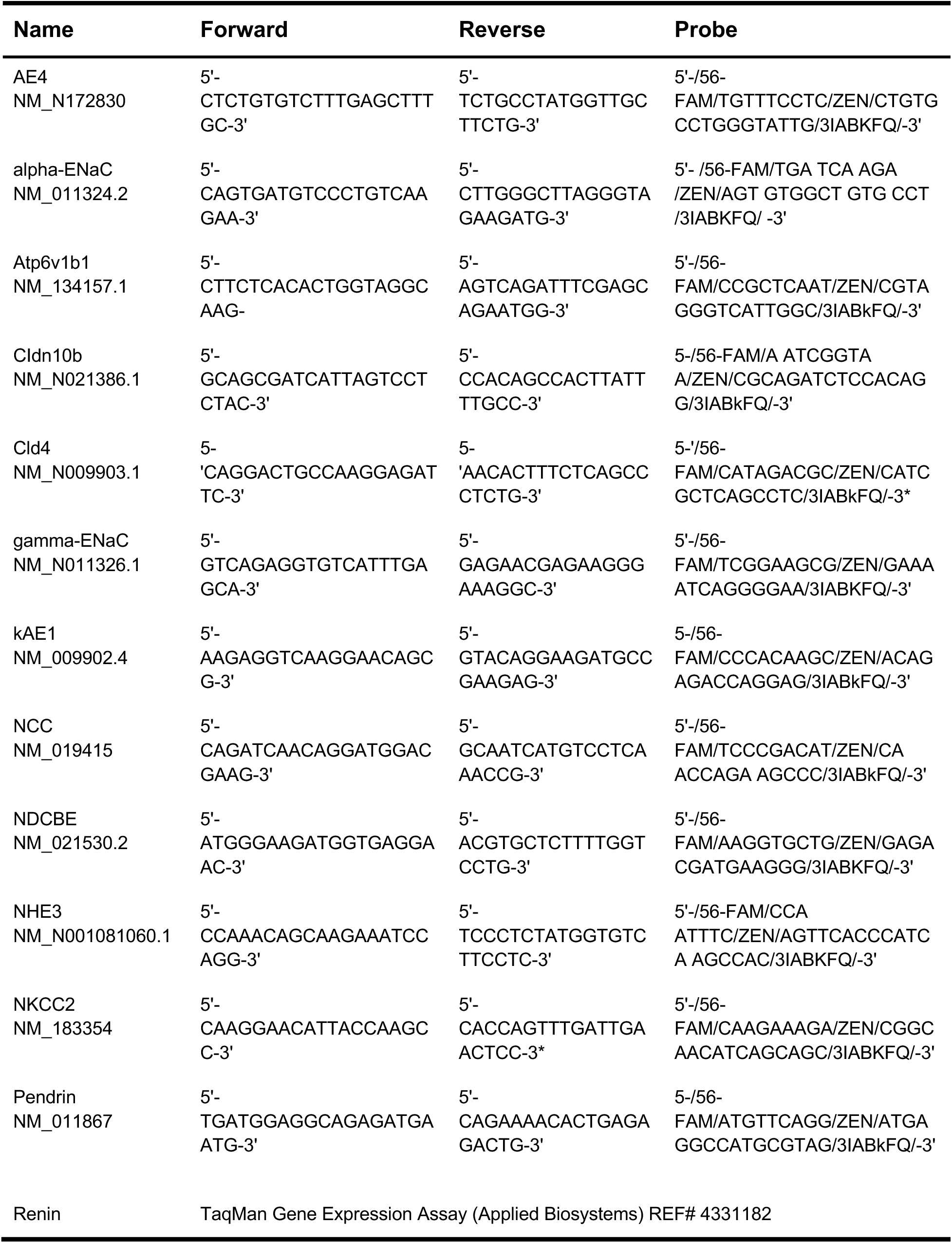
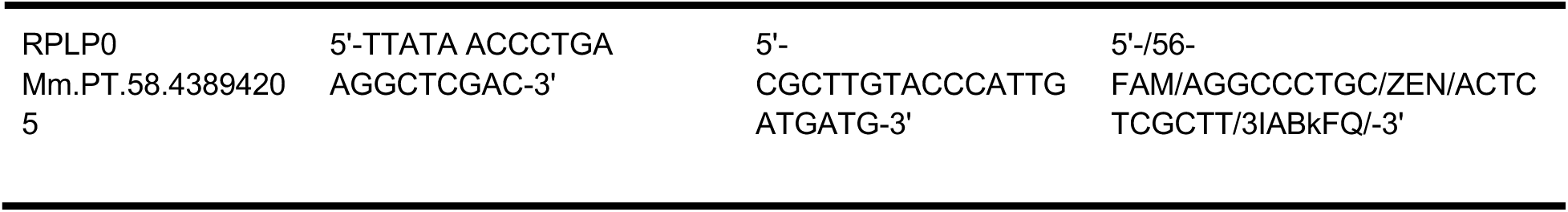
Primers and Probes Used for qPCR on Mouse Genes.

**Supplementary Table 2:** Antibodies Used for Immunoblotting.

| Name | Reference | Reference | Dilution |
| --- | --- | --- | --- |
| Rb anti-Claudin-10 | Primary | Antibodies.com | 1:1000 |
| Rb anti-Claudin-4 | Primary | Invitrogen | 1:1000 |
| Rb anti-ENaC<br>gamma | Primary | StressMarq<br>Biosciences Inc | 1:5000 |
| Direct-Blot HRP<br>anti-β Actin | Primary | BioLegend | 1:10000 |

**Supplementary Table 3.** Baseline Plasma Characterization of WT and L919X Ae1 KI/KI Mice.

| Parameter | Units | WT (n=10) | L919X AE1 KI/KI (=10) | p-value |
| --- | --- | --- | --- | --- |
| Na<sup>+</sup> | mM | 147.5 ± 0.563 | 146.9 ± 0.623 | 0.4327 |
| Cl<sup>-</sup> | mM | 118.8 ± 0.964 | 119.6 ± 1.536 | 0.6644 |
| K<sup>+</sup> | mM | 4.633 ± 0.167 | 4.610 ± 0.196 | 0.7029 |
| HCO<sub>3</sub><sup>-</sup> | mM | 18.38 ± 0.563 | 19.95 ± 0.511 | 0.0930 |
| pH |  | 7.289 ± 0.002 | 7.249 ± 0.009 | 0.1186 |
| TCO<sub>2</sub> | mmHg | 19.50 ± 0.749 | 21.30 ± 0.517 | 0.0636 |
| BUN | mg/dl | 22.70 ± 1.382 | 19.80 ± 0.987 | 0.1050 |
| Glucose | mg/dl | 181 ± 13.11 | 198.3 ± 10.58 | 0.3181 |
| HCT | % | 36.80 ± 0.554 | 36 ± 0.789 | 0.4174 |
| pCO<sub>2</sub> | mmHg | 38.99 ± 2.856 | 45.75 ± 1.595 | 0.0525 |
| AnGAP | mM | 14.6 ± 0.618 | 13.22 ± 1.325 | 0.1197 |
| Hb | g/dl | 12.53 ± 0.188 | 12.23 ± 0.269 | 0.3729 |
| BE<sub>ecf</sub> | mM | -8.100 ± 0.781 | -7.100 ± 0.504 | 0.2963 |

*Data is presented as mean ± SEM. Student's T-Test or Mann-Whitney test used where appropriate.*

**Supplementary Table 4.** Baseline Plasma Characterization of WT and R607H Ae1 KI/KI Mice.

| Parameter | Units | WT (n=11) | R607H AE1 KI/KI (n=9) | p-value |
| --- | --- | --- | --- | --- |
| Na<sup>+</sup> | mM | 146.4 ± 0.472 | 147.1 ± 0.586 | 0.3148 |
| Cl<sup>-</sup> | mM | 118.2 ± 1.110 | 118.5 ± 0.619 | 0.1409 |
| **K<sup>+</sup>** | **mM** | **4.5 ± 0.142** | **4.920 ± 0.077\*** | **0.0207** |
| HCO<sub>3</sub><sup>-</sup> | mM | 18.66 ± 0.627 | 19.07 ± 0.463 | 0.6142 |
| pH |  | 7.256 ± 0.014 | 7.258 ± 0.013 | 0.9102 |
| TCO<sub>2</sub> | mmHg | 19.91 ± 0.653 | 20.20 ± 0.533 | 0.7370 |
| **BUN** | **mg/dl** | **21.09 ± 1.351** | **23.60 ± 0.921\*** | **0.0438** |
| Glucose | mg/dl | 196.3 ± 7.065 | 197.3 ± 12.31 | 0.9417 |
| HCT | % | 37.73 ± 1.221 | 38.60 ± 0.340 | 0.5178 |
| pCO<sub>2</sub> | mmHg | 42.05 ± 1.467 | 42.72 ± 1.410 | 0.7417 |
| AnGAP | mM | 13.27 ± 1.538 | 14.60 ± 0.600 | 0.9863 |
| Hb | g/dl | 12.82 ± 0.414 | 13.14 ± 0.117 | 0.4826 |
| BE<sub>ecf</sub> | mM | -8.545 ± 0.731 | -8.100 ± 0.586 | 0.5666 |

*Data is presented as mean ± SEM. Student's T-Test or Mann-Whitney test used where appropriate. **Significant data is bolded.***

**Supplementary Table 5.** Plasma Characterization of WT and L919X Ae1 KI/KI Mice Following 24 hours of Salt Restriction.

| Parameter | Units | WT (n=10) | L919X AE1 KI/KI (n=10) | p-value |
| --- | --- | --- | --- | --- |
| Na<sup>+</sup> | mM | 147.9 ± 0.690 | 147.7 ± 0.578 | 0.8268 |
| Cl<sup>-</sup> | mM | 117.6 ± 0.897 | 120.9 ± 1.329 | 0.0543 |
| K<sup>+</sup> | mM | 4.980 ± 0.158 | 5.180 ± 0.464 | 0.5661 |
| HCO<sub>3</sub><sup>-</sup> | mM | 20.59 ± 0.613 | 20.40 ± 0.825 | 0.8554 |
| pH |  | 7.240 ± 0.021 | 7.248 ± 0.033 | 0.8469 |
| TCO<sub>2</sub> | mmHg | 22.20 ± 0.593 | 21.80 ± 0.854 | 0.7048 |
| BUN | mg/dl | 30.10 ± 4.365 | 23.30 ± 0.989 | 0.1249 |
| Glucose | mg/dl | 164.5 ± 12.69 | 157.1 ± 9.719 | 0.6490 |
| HCT | % | 40.70 ± 0.667 | 38.90 ± 0.690 | 0.1152 |
| pCO<sub>2</sub> | mmHg | 47.95 ± 1.127 | 47.74 ± 3.411 | 0.9540 |
| AnGAP | mM | 14.90 ± 0.795 | 11.89 ± 1.369 | 0.0677 |
| Hb | g/dl | 13.84 ± 0.225 | 13.22 ± 0.237 | 0.0741 |
| BEecf | mM | -6.900 ± 0.971 | -6.800 ± 1.073 | 0.9457 |

*Data is presented as mean ± SEM. Student's T-Test or Mann-Whitney test used where appropriate. **Significant data is bolded.***

**Supplementary Table 6.** Plasma Characterization of WT and R607H Ae1 KI/KI Mice Following 24 hours of Salt Restriction.

| Parameter | Units | WT (n=12) | R607H Ae1 KI/KI (n=14) | p-value |
| --- | --- | --- | --- | --- |
| Na<sup>+</sup> | mM | 146.3 ± 0.890 | 148.1 ± 0.275 | 0.8166 |
| Cl<sup>-</sup> | mM | 121.5 ± 2.062 | 122.9 ± 0.769 | 0.5196 |
| **K<sup>+</sup>** | **mM** | **4.475 ± 0.100** | **4.843 ± 0.100** | **0.0162** |
| HCO<sub>3</sub><sup>-</sup> | mM | 19.04 ± 0.965 | 18.39 ± 0.540 | 0.5437 |
| pH |  | 7.203 ± 0.021 | 7.215 ± 0.014 | 0.6389 |
| TCO<sub>2</sub> | mmHg | 20.50 ± 1.004 | 19.86 ± 0.592 | 0.9305 |
| BUN | mg/dl | 23.75 ± 1.473 | 25.00 ± 1.267 | 0.5236 |
| Glucose | mg/dl | 154.5 ± 11.59 | 182.6 ± 9.245 | 0.0667 |
| HCT | % | 37.50 ± 2.183 | 37.86 ± 0.710 | 0.1096 |
| pCO<sub>2</sub> | mmHg | 48.33 ± 2.356 | 45.75 ± 1.901 | 0.3983 |
| AnGAP | mM | 10.25 ± 2.394 | 11.64 ± 0.608 | 0.1453 |
| Hb | g/dl | 12.76 ± 0.741 | 12.86 ± 0.246 | 0.0994 |
| BE<sub>ecf</sub> | mM | -9.000 ± 1.206 | -9.357 ± 0.608 | 0.7847 |

*Data is presented as mean ± SEM. Student's T-Test or Mann-Whitney test used where appropriate. **Significant data is bolded.***

**Supplementary Table 7.** Plasma Characterization of WT and L919X Ae1 KI/KI Mice Following Salt Depleted Acid Load.

| Parameter | Units | WT (n=14) | L919X AE1 KI/KI (n=14) | p-value |
| --- | --- | --- | --- | --- |
| **Na<sup>+</sup>** | **mM** | **149.4 ± 0.644** | **156.4 ± 1.152** | **< 0.0001** |
| **Cl<sup>-</sup>** | **mM** | **126.9 ± 2.087** | **135.1 ± 0.900** | **0.0140** |
| K<sup>+</sup> | mM | 4.777 ± 0.154 | 4.669 ± 0.170 | 0.6430 |
| HCO<sub>3</sub><sup>-</sup> | mM | 15.63 ± 0.940 | 13.88 ± 0.470 | 0.1164 |
| pH |  | 7.170 ± 0.030 | 7.131 ± 0.014 | 0.0683 |
| **TCO<sub>2</sub>** | **mmHg** | **17.62 ± 0.789** | **15.15 ± 0.504** | **0.0147** |
| **BUN** | **mg/dl** | **23.75 ± 0.954** | **33.62 ± 1.723** | **< 0.0001** |
| Glucose | mg/dl | 148.9 ± 12.93 | 131.92 ± 5.054 | 0.2452 |
| HCT | % | 37.38 ± 1.274 | 37.46 ± 1.264 | 0.9713 |
| pCO<sub>2</sub> | mmHg | 42.84 ± 2.581 | 41.72 ± 1.632 | 0.7216 |
| AnGAP | mM | 13.33 ± 1.694 | 13.09 ± 0.948 | 0.9042 |
| Hb | g/dl | 12.70 ± 0.432 | 12.74 ± 0.430 | 0.9713 |
| BE<sub>ecf</sub> | mM | -12.93 ± 1.294 | -15.31 ± 0.603 | 0.1169 |

*Data is presented as mean ± SEM. Student's T-Test or Mann-Whitney test used where appropriate. **Significant data is bolded.***

**Supplementary Table 8.** Plasma Characterization of WT and R607H Ae1 KI/KI Mice Following Salt Depleted Acid Load.

| Parameter | Units | WT (n=14) | R607H AE1 KI/KI<br>(n = 18) | p-value |
| --- | --- | --- | --- | --- |
| Na<sup>+</sup> | mM | **145.2 ± 0.882** | **156.4 ± 0.903** | **< 0.0001** |
| Cl<sup>-</sup> | mM | **126.8 ± 1.959** | **138.6 ± 0.536** | **< 0.0001** |
| K<sup>+</sup> | mM | 4.557 ± 0.229 | 4.406 ± 0.185 | 0.6069 |
| HCO<sub>3</sub><sup>-</sup> | mM | **14.15 ± 0.921** | **11.64 ± 0.775** | **0.0448** |
| pH |  | **7.192 ± 0.025** | **7.085 ± 0.020** | **0.0016** |
| TCO<sub>2</sub> | mmHg | 15.29 ± 0.963 | 12.83 ± 0.797 | 0.0572 |
| BUN | mg/dl | **22.50 ± 1.725** | **34.17 ± 3.618** | **0.0026** |
| Glucose | mg/dl | **163.2 ± 11.93** | **118.8 ± 7.284** | **0.0024** |
| HCT | % | 34.00 ± 2.176 | 32.56 ± 2.297 | 0.8434 |
| pCO<sub>2</sub> | mmHg | 46.21 ± 1.429 | 38.56 ± 2.400 | 0.4397 |
| AnGAP | mM | 10.08 ± 2.065 | 13.43 ± 1.462 | 0.6004 |
| Hb | g/dl | **11.56 ± 0.739** | **13.55 ± 0.189** | **0.0489** |
| BEecf | mM | **-14.07 ± 1.273** | **-18.22 ± 0.967** | **0.0129** |

*Data is presented as mean ± SEM. Student's T-Test or Mann-Whitney test used where appropriate. **Significant data is bolded.***

