## Supplementary Material for "Urinary Sodium Wasting and Disrupted Collecting Duct Function in Mice with dRTA-Causing *SLC4A1* Mutations"

Priyanka Mungara[1,2], Kristina MacNaughton[1,2], AKM Shahid Ullah[1,2], Grace Essuman[1,2], Forough Chelangarimiyandoab[1,2], Rizwan Mumtaz[4], J. Christopher Hennings[4], Christian A. Hübner[4,5], Dominique Eladari[6,7,8], R. Todd Alexander[1,2,3], Emmanuelle Cordat[1,2*]

### Materials and Methods

*Animals.* In Germany, all animal experiments were approved by the Thüringer Landesamt für Verbraucherschutz (TLV) under the license 02-016/14. In Canada, all animal protocols were approved by the University of Alberta's Animal Care and Use Committee (AUP #1277) and were in accordance with the National and institutional Animal care guidelines. At baseline, mice were maintained on a 12-h light/dark cycle with drinking water and standard rodent chow (PicoLab Mouse Diet 20 # 5058) available *ad libitum*. R607H KI mice (129/SvJ strain) have been previously published and their generation described[32].

*Generation of the L919X KI mice*. L919X KI mice were generated by homologous recombination similar to Ae1 R607H KI mice[33]. Briefly, a 14.7 kb genomic DNA fragment (GRCm38/mm10; chr11:102,341,468-102,356,137) from a lambda phage library including exons 14-20 of the Slc4a1 gene was cloned into the pKO-DTA Scrambler vector (Lexicon Genetics) containing a phosphoglycerate kinase (Pgk) promoter driven diphteria toxin A cassette. A neomycin resistance cassette flanked by loxP sites and an 5´ EcoRV site were introduced at the BspHI restriction site in the genomic Ae1 sequence. Site-directed mutagenesis was used to generate the L919X mutation in the adjacent exon 20. The linearized targeting vector was electroporated into R1 mouse embryonic stem cells. Clones resistant to Neomycin were analyzed by Southern blot on EcoRV-digested genomic DNA with an external probe (GRCm38/mm10; chr11:102,356,140-102,356,838) labeling a 9.1 kb wild-type or 6.1 kb knock-in fragment (**Figure 1 A**). Correctly targeted ES-clones were injected into C57BL6 blastocysts to obtain chimeric mice. Chimeric mice with germ line transfer were bred with a Cre-Deleter mouse strain to excise the Neomycin resistance cassette. Heterozygous KI mice were backcrossed for at least 5 generations with C57BL/6 mice before WT and homozygous KI mice (KI/KI) were bred for experiments. Mice were genotyped by polymerase chain reaction (PCR) of ear notch biopsies using the following primers for the R607H mice: 5′-TAG CTC CTT CTA CCC CAC CCA-3′ and 5′-CCA GAG GTA CAT GGT AAA ACA TTG TC-3′, as previously published [32] and the following primers for the L919X mice: 5′-GCT TCC GTC TTG GTC TGC TGT G-3′ and 5′-TGG ACA AGC CCT GCT GTC CCT A-3', detecting a 301 bp band for WT allele and a 444 bp for the KI allele.

*Metabolic Cage Experimental Design and Diets.* For each diet, 2 - 4 months old homozygous KI (R607H or L919X) mice and WT littermates (equal number male and female) were placed in metabolic cages (Tecniplast) for 72 hours to acclimate and were fed a standard rodent chow and water (PicoLab Mouse Diet 20 # 5058), available *ad libitum*. Following this period, mice were fed one of two diets:

- Protocol 1 "Salt Depletion": R607H KI/KI, L919X KI/KI mice, and WT littermates were fed a salt restricted diet (0.01 - 0.02 % sodium, 0.07 % chloride, Teklad Custom Diet # TD.08290) and given regular water *ad libitum* for 24 hours.
- Protocol 2 "Salt Depletion with Acid Load": R607H KI/KI, L919X KI/KI, and WT littermates were fed a low salt diet for 8 days (# TD.08290). On day 9, the mice continued this salt restriction but were provided with acid in their drinking water (0.28 M $NH_4Cl$ with 0.5 % sucrose), available *ad libitum* for 6 additional days. Fresh acid water was replaced every day. Total experimental period was 14 days.

Body weight, chow and water consumption, urine (under mineral oil), and feces measurements were collected every 24 hours. On the final experimental day of *Protocol 1* & *2*, mice were anesthetized with a 50 mg/kg dose of pentobarbital. Blood was collected in lithium heparin-coated tubes and centrifuged for 10 minutes at 4°C to collect serum which was then stored at -80 °C. After PBS/heparin perfusion, whole kidneys were dissected into 4 halves and each half was processed for either RNA extraction, immunoblot, immunofluorescence, or immunohistochemistry, and then stored at -80 °C.

*Urine and Serum Analysis.* Freshly collected blood was analyzed for electrolytes ($Na^+$, $K^+$, and $Cl^-$), glucose, urea nitrogen (BUN), hematocrit (Hct) and hemoglobin (Hgb), and pH using an i-STAT1 Analyzer (Abaxis) with an i-STAT Chem8+ cartridge chip (Abbott Laboratories). Urine pH was measured using pH microelectrode (PerpHect Ross Micro Combination pH electrode, ThermoScientific) attached to a Accumet AR10 pH meter (Fischer Scientific). Urine osmolality was measured with the Advanced Instruments Osmo1 Single-Sample Micro-Osmometer (Thermo Fisher Scientific), diluting samples at 1:10 or 1:100 with filtered $ddH_2O$**.** Urine electrolytes were measured by ion chromatography (Dionex Aquion Ion Chromatography System, Thermo Fisher Scientific Inc.) with an autosampler. Urine samples were diluted 1:100 or 1:250 with $ddH_2O$ and a 4.5 mM $Na_2CO_3$/1.5 mM $NaHCO_3$ in $ddH_2O$ solution was used for anion eluent, and a 20 mM methanesulfonic acid in $ddH_2O$ solution for cation eluent composition[36]. Urine creatinine was measured using a parameter creatinine kit (R&D systems) or with ion chromatography. Chromeleon 7 Chromatography Data System software (Thermo Scientific) was used to analyze results. Urine ions were normalized to urinary creatinine concentration.

*RNA isolation and Reverse Transcription Quantitative PCR.* Immediately after collection, a half kidney was incubated in RNAlater (ThermoFisher) for 8 hours at room temperature before storing in -80°C. Total RNA was extracted using TRIzol reagent (Invitrogen) as previously described [37]. RNA quantification was performed using NanoDrop (NanoDrop 2000C, Thermo Fisher Scientific). 2 µg of cDNA was reverse transcribed with SuperScript™ II (Invitrogen) as per the manufacturer's instructions. The cDNA was pooled to create a serially diluted standard curve. Real-time (RT) quantitative PCR was performed in triplicate for each cDNA sample using TaqMan PCR master mix in a QuantStudio 6 Pro RT PCR system (ThermoFisher Scientific). Samples were quantified using the $2^{-\Delta\Delta Ct}$ method[38]. Expression levels of mRNA from specific genes were normalized to housekeeping ribosomal protein lateral stalk subunit P0 (RPLP0) gene. Primers for the genes of interest are listed in **Supplementary Table 1.**

*Protein Extraction and Immunoblot.* A quarter of a freshly dissected kidney was decapsulated and mechanically homogenized in ice-cold lysis buffer (0.3 M Sucrose, 25 mM Imidazole, 1mM EDTA, 8.5 µM Leupeptin, 1 mM PMSF) [39]**,** vortexed every 15 mins over 1 hour, and centrifuged at 4°C, 14000 rpm for 15 minutes. Protein concentration was measured using a Bicinchoninic Acid Protein Assay. Immunoblot experiments were performed with 8 %, 10 %, or 12 % SDS-PAGE gels, transferred to a PVDF membrane, and blocked with 3 % milk in TBST (5 mM Tris base, 15 mM NaCl, 0.1% Tween 20). Membranes were incubated in primary antibodies overnight at 4°C followed by incubation in secondary antibodies conjugated to horseradish peroxidase. The antibodies used are listed in **Supplementary Table 2**. Proteins were visualized using the Clarity Western ECL kit (Bio-Rad) and images were captured by ChemiDoc touch imaging systems (Bio-Rad). Quantification and densitometric analysis were performed by ImageJ (National Institutes of Health, USA).

*Immunostaining*. Freshly collected kidneys retrogradely perfused with PFA (4 %) were flash frozen in liquid nitrogen cooled isopentane. Cryosections (4 µm thickness) were prepared and either subjected to Masson-Goldner stain, Von Kossa staining or further processed for immunostaining as previously described[40].

*Statistical Analysis.* Statistical analysis was completed using GraphPad Prism software (Ver 7. 0e). Normality was verified for all data sets and outliers (as determined by Prism) were removed. Analysis was performed using unpaired Student's t-test or Mann-Whitney test, or Two-Way ANOVA where appropriate. A p-value of less than 0.05 was statistically significant. All data are presented as mean ± SEM.

**Supplementary Figure legends**

**Supplementary Figure 1**: Von Kossa (A) and Masson Trichrome (B) staining of WT, heterozygous (L919X KI) or homozygous (L919X KI/KI) mouse cortical or inner stripe of outer medullary kidney sections.

**Supplementary Figure 2**: After a 24h salt-depleted diet, both R607H and L919X KI/KI mice continue to produce an alkaline urine compared to WT littermates. (**A**) urinary pH at baseline (left) and after 18 h of diet (right) in WT or Ae1 L919X KI/KI mice. Urine osmolality (**B**), urinary sodium/creatinine ratio (**C**), urinary chloride/creatinine ratio (**D**) and urinary potassium/creatinine ratio (**E**) over the course of the 24h experiment in WT or L919X KI/KI mice. (**F**) urinary pH at baseline (left) and after 18 h of diet (right) in WT or Ae1 R607H KI/KI mice. Urine osmolality (**G**), urinary sodium/creatinine ratio (**H**), urinary chloride/creatinine ratio (**I**) and urinary potassium/creatinine ratio (**J**) over the course of the 24h experiment in WT or L919X KI/KI mice. Error bars correspond to means ± SEM, **P < 0.01, ***P < 0.001, ****P < 0.0001 using Student's t-test or Mann-Whitney test, or two-way ANOVA with Tukey's multiple comparison test.

**Supplementary Figure 3**: Comparison of gene expression and protein abundance of markers of the IC, PC and TAL in L919X KI/KI mice or WT littermates. Gene expression of A-IC markers kAE1 (**A**), B1-ATPase (**B**) and protein expression of B1-ATPase (**C**) with a representative immunoblot image (right). Gene expression of B-IC markers pendrin (**D**), NDCBE (**E**) and AE4 (**F**). Gene expression of PC markers ROMK (**G**), gamma-ENaC (**H)** and gamma-ENaC protein abundance (**I)** with representative immunoblot image (right). Gene expression of TAL and DCT markers NCC (**J**), NHE3 (**K**) and NKCC2 (**L**). Error bars correspond to means ± SEM, **P < 0.01, ***P < 0.001, ****P < 0.0001 using Student's t-test.

**Supplementary Figure 4**: Comparison of gene expression and protein abundance of markers of the IC, PC and TAL in R607H KI/KI mice or WT littermates. Gene expression of A-IC markers kAE1 (**A**), B1-ATPase (**B**) and protein expression of B1-ATPase (**C**) with a representative immunoblot image (right). Gene expression of B-IC markers pendrin (**D**), NDCBE (**E**) and AE4 (**F**). Gene expression of PC markers ROMK (**G**), gamma-ENaC (**H)** and gamma-ENaC protein abundance (**I)** with representative immunoblot image (right). Gene expression of TAL and DCT markers NCC (**J**), NHE3 (**K**) and NKCC2 (**L**). Error bars correspond to means ± SEM, *P < 0.05, **P < 0.01, ***P < 0.001, ****P < 0.0001 using Student's t-test.

A

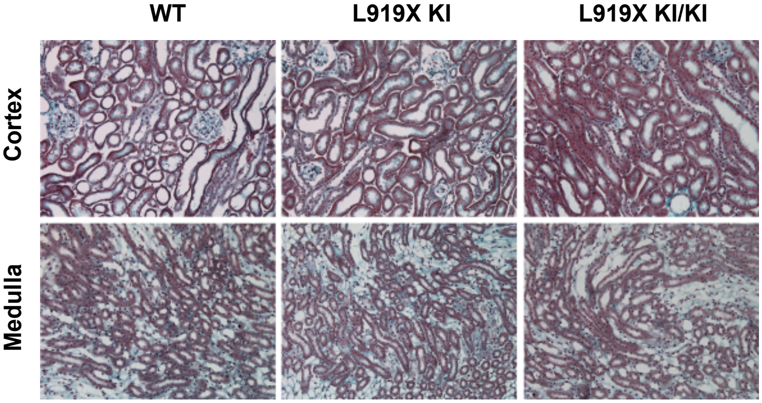


B

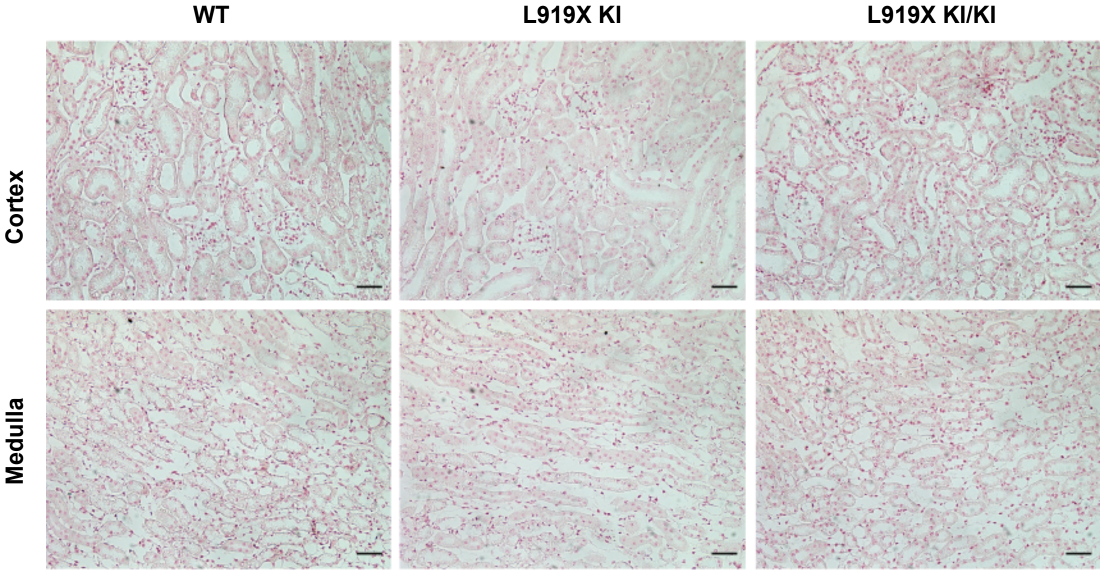


Supplementary Figure 1 Mungara et al.

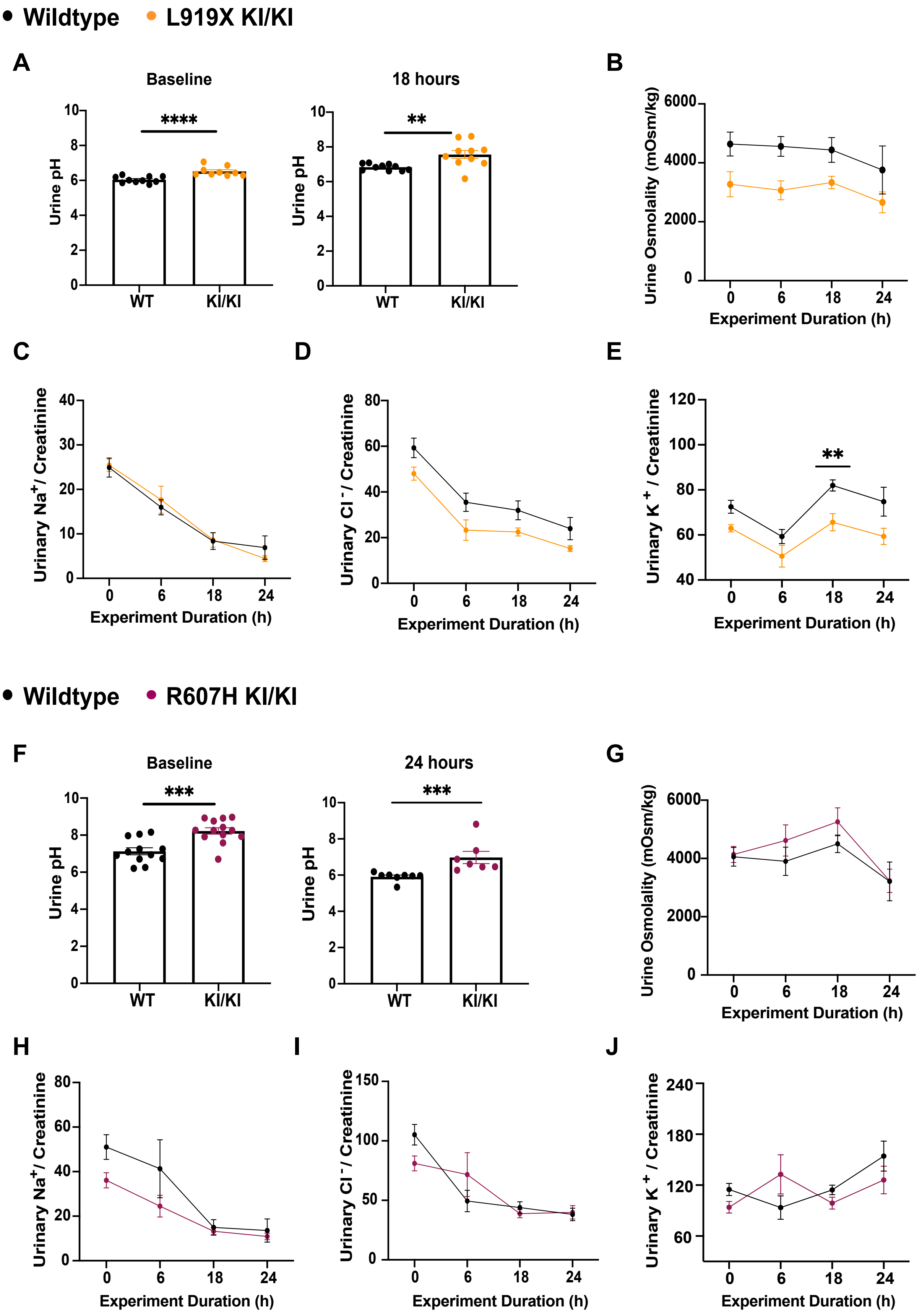


Supplementary Figure 2 Mungara et al.

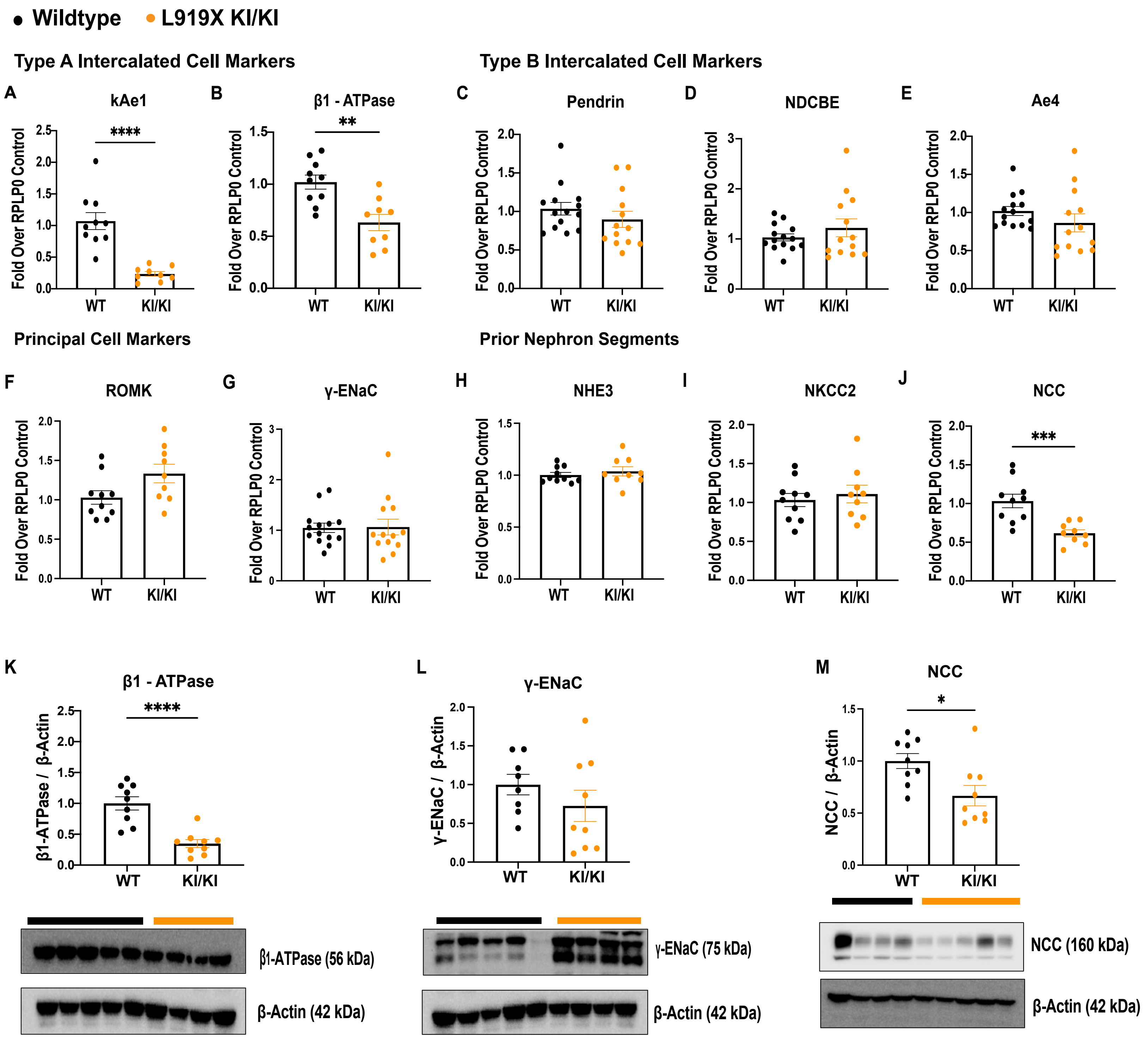


Supplementary Figure 3 Mungara et al.

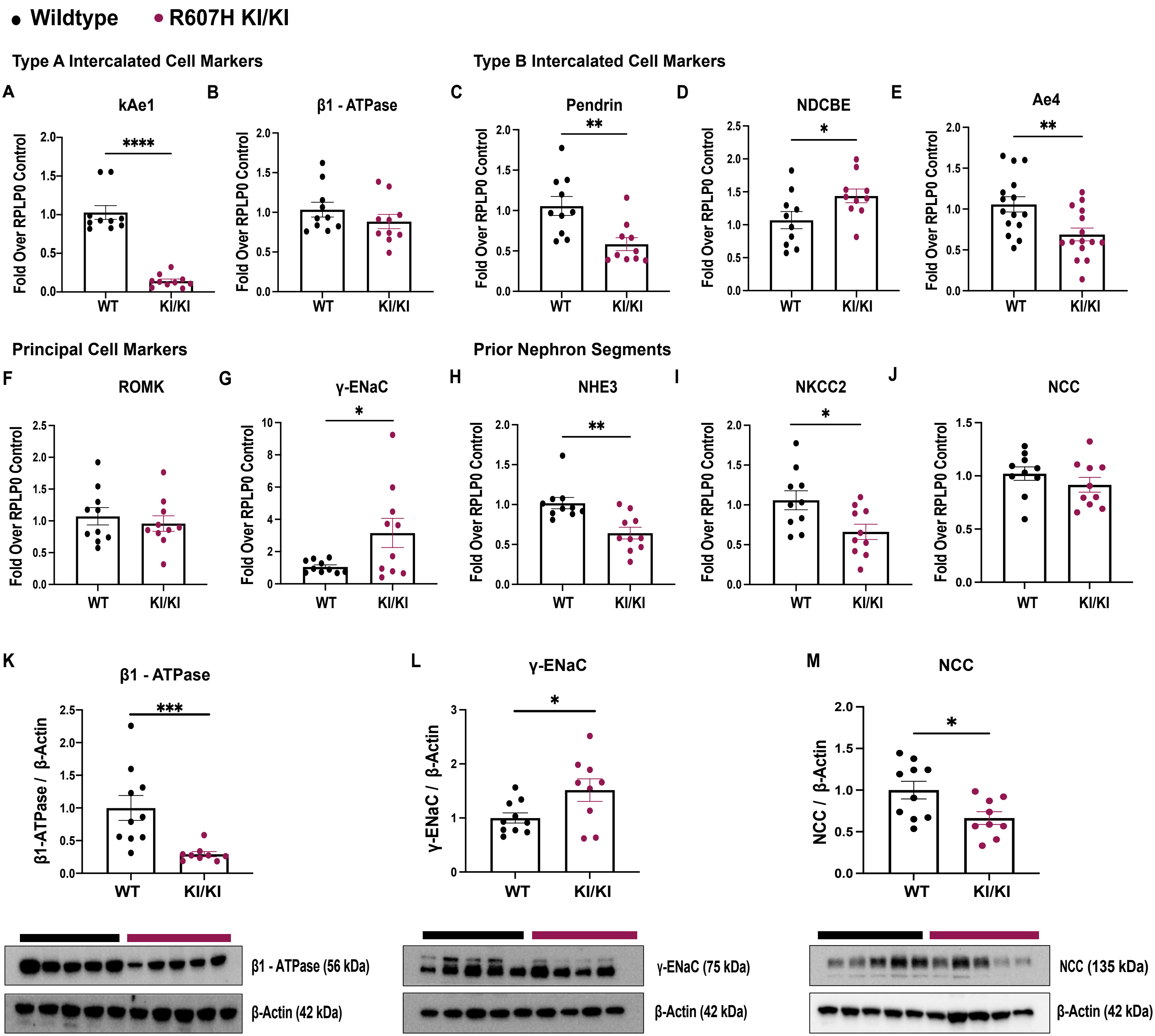


Supplementary Figure 4 Mungara et al.
